# Function of cell adhesion molecules in differentiation of ray sensory neurons in *C. elegans*

**DOI:** 10.1101/2022.09.12.507584

**Authors:** Naoko Sakai, Peter Sun, Byunghyuk Kim, Scott W. Emmons

**Affiliations:** Department of Genetics, Albert Einstein College of Medicine, United States; Department of Physiology, Tokyo Women’s Medical University, Tokyo, Japan; Department of Life Science, Dongguk University, South Korea

## Abstract

For proper functioning of the nervous system, it is crucial that neurons find their appropriate partners and build the correct neural connection patterns. Although cell adhesion molecules (CAMs) have been studied for many years as essential players in neural connections, we have yet to unravel the code by which CAMs encode synaptic specificity. We analyzed the effects of mutations in CAM genes on the morphology and synapses of a set of sensory neurons in the *C. elegans* male tail. B-type ray sensory neurons express ten genes encoding CAMs. We examined the effect on axon trajectory and localization of presynaptic components in viable mutants of nine of these. We found axon trajectory defects in mutants of UNC-40/DCC, SAX-3/ROBO, and FMI-1/Flamingo/Celsr1. In none of the mutants was presence of presynaptic components in axons lost, and in several the level appeared to increase, suggesting possible accumulation. B-type sensory neurons fasciculate with a second type of ray sensory neuron, the A-type, in axon commissures. We found cell non-autonomous effects consistent with each promoting the trajectory of the other. Overall, single and multiple mutants of CAM genes had limited effects on ray neuron trajectories and accumulation of synaptic components.

## Introduction

Since the proposal that the pattern of neural connections is controlled by individual identification tags on the cell surfaces of neurons (Sperry, 1963), a number of cell adhesion molecules (CAMs) that regulate axon outgrowth and synapse formation have been identified through forward genetic and other methods (Hedgecock et al., 1990; Zallen et al., 1998; Yamagata et al., 2002; Shen and Bargmann, 2003; Salzberg et al., 2013; Carrillo et al., 2015; Kurshan et al., 2018; Philbrook et al., 2018; Kurshan and Shen, 2019; Taylor et al., 2021). Despite efforts in the last few decades, we have yet to break the code by which cell adhesion molecules help neurons find their appropriate partners. This is due in part to the redundant and pleiotropic functions of CAMs. There is limited information on expression patterns of CAMs in circuits of known connectivity, which has limited us in studying these parallel functions.

The nematode *C. elegans* has been a well-studied model organism for neuronal development because of its invariant developmental pattern and fully reconstructed neural and synaptic connectivity (White et al., 1986; Cook et al., 2019). Because of this feature, *C. elegans* has played an important role in studies of the relationship between CAMs and development (Yamagata et al., 2002; Shen et al., 2004; Salzberg et al., 2013; Kim and Emmons, 2017; Shah et al., 2017; Lázaro-Peña et al., 2018; Philbrook et al., 2018; Taylor et al., 2021) In this study, we focused on a set of male-specific sensory neurons to investigate the relationship between cell adhesion molecules, neurite trajectory, and synapse formation.

The *C. elegans* male has a set of nine bilateral pairs of sensory structures known as rays projecting from the tail embedded within a cuticular fan (Fig. 1*A*). Each ray has a similar structure consisting of the sensory endings of two sensory neurons of two types designated A-type and B-type (Sulston et al., 1980). In addition, each of the nine ray pairs has a distinct identity affecting the neurotransmitters, synaptic targets, and functions of the constituent neurons (Baird et al., 1991; Chow and Emmons, 1994; Lints et al., 2004). As part of a large cohort of male-specific neurons and muscles, the ray sensory neurons are born during the L3 larval stage and differentiate through the L4 and into young adulthood (Sulston et al., 1980). Their cell bodies are located in a pair of bilateral lumbar ganglia. To reach their synaptic targets, ray neurons extend axons through circumferential commissures and into the pre-anal ganglion, where their growing processes branch considerably and synapse onto a number of post-synaptic targets (Fig. 1*A,B*). In this study, we focus on the role of CAMs in determining the trajectories and synapse formation of the B-type ray neurons (RnB, n = 1-9, designating the nine rays).

**Fig. 1.**
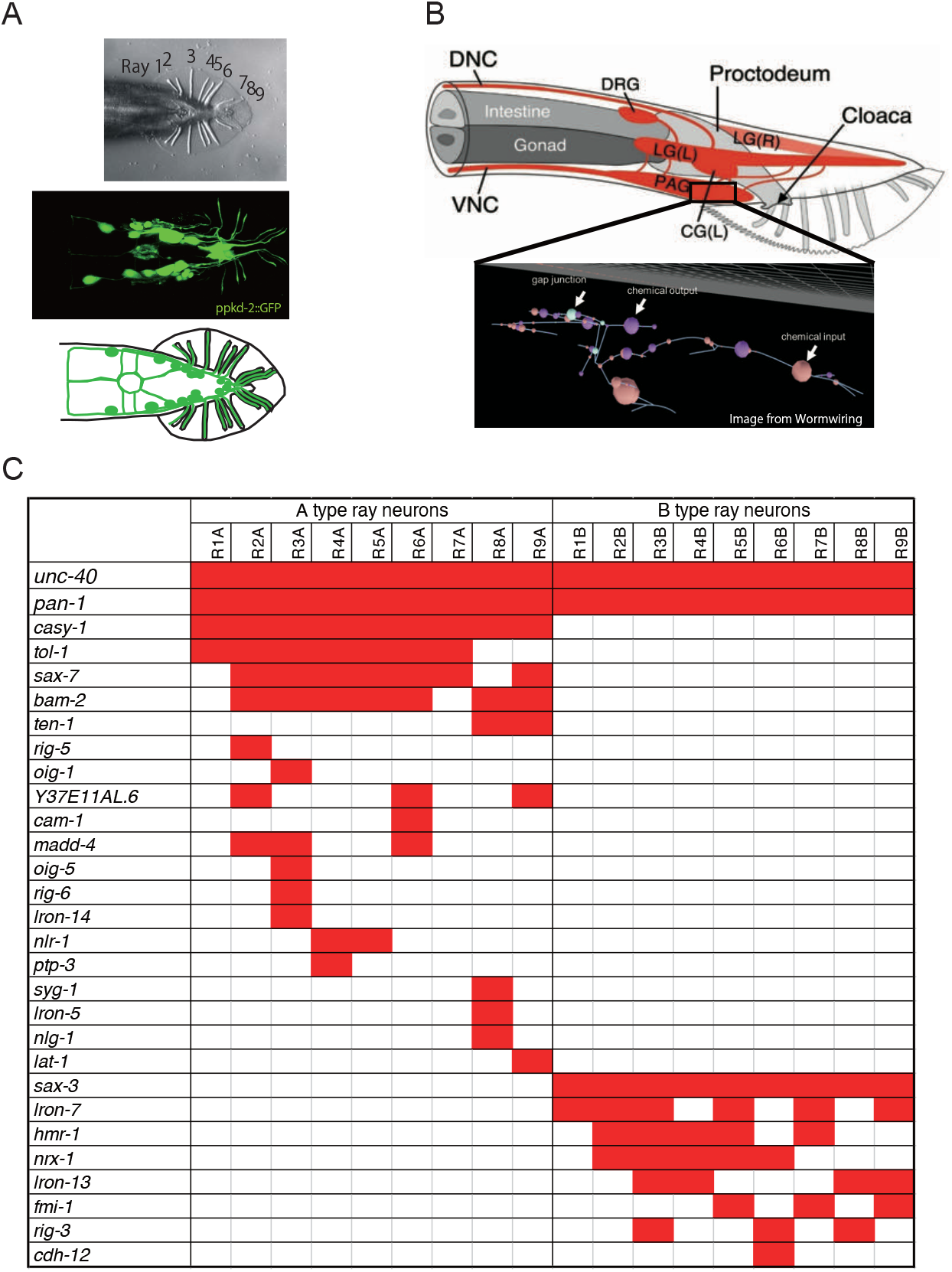
CAMs expressing in Ray neurons. (A) A Nomarski image of Ventral view of male tail (top), a Fluorescent image of RnB neuron processes marked by pkd-2^prom^::GFP (middle), and schematic image of male ray structure and RnB processes (bottom). (B) Schematic image of lateral view of male tail (top) and chemical and electrical junctions of R2BL in the pre anal ganglion (bottom). (C) Expression of transcriptional reporters for CAMs in the RnA and RnB neurons. One hundred CAMs genes have been examined and the genes having expression in the ray neurons are shown as red boxes.

We have determined the expression patterns of the cell adhesion genes encoded in the *C. elegans* genome across the *C. elegans* posterior nervous system (in preparation). In our dataset, the B-type ray neurons collectively express 10 different CAMs, while the A-types altogether express 21 (Fig 1*C*). The pattern is left/right symmetric — in each case, the left/right homologous neurons had the same expression, and it is ray and neuron-type specific — only two of the genes, *unc-40* and *pan-1*, are expressed in both A and B-types. In this study, we examined the effects of mutations in the B-type-expressed genes as well as several of the A-type expressed genes on the ray B-type axon trajectories and synapse formation, using reporters that allow us to simultaneously score both the neurite trajectory and the localization of pre-synaptic components. In a previous forward genetic screen, it was found that mutations in *unc-40/DCC* and *sax-3/ROBO* prevented outgrowth of the B-type axons through the circumferential commissures (Jia and Emmons, 2006). In a second study, B-type neuron synapse formation was affected by mutations in CAMs neurexin and neuroligin as well as by mutation in a modification enzyme that affects the sulfation pattern of sugar residues on the matrix protein glypican (Lázaro-Peña et al., 2018). Here we demonstrate effects of mutations in *fmi-1*/flamingo and a presumptively non-autonomous effect of mutation in a gene expressed in A-neurons, *sax-7*/L1CAM, on B-neuron trajectory. Overall, however, in our nearly complete survey, we found limited effects of both single and multiple mutations in most of the expressed CAM genes.

## Materials and methods

### Animal Maintenance

All animals were maintained using standard methods (Brenner, 1974). Briefly, all animals were grown on NGM plates seeded with E. coli OP50 at 20°C. All strains contain the *<u>him-5</u>(<u>e1490</u>)* mutation to increase the male population(Broverman and Meneely, 1994) unless otherwise specified. For all strain information, see the supplementary file.

### Statistical analyses

All data was analyzed by Fisher’s exact test or multiple comparison test using Graph-pad Prism 9.0 statistical software (RRID:SCR_002798).

### Plasmid constructions and germ-line transformation

Plasmid constructs were generated using the In-Fusion HD Cloning Kit (Takara-bio). To construct pkd-2prom:: 3xnovoGFP-CLA-1, we amplified pkd-2 promoter with the primers as listed and fused into PK085 plasmid (unc-129^prom^::3xnovoGFP-CLA1, kindly provided by P. Kurshan) digested with xbaI and sphI sites.

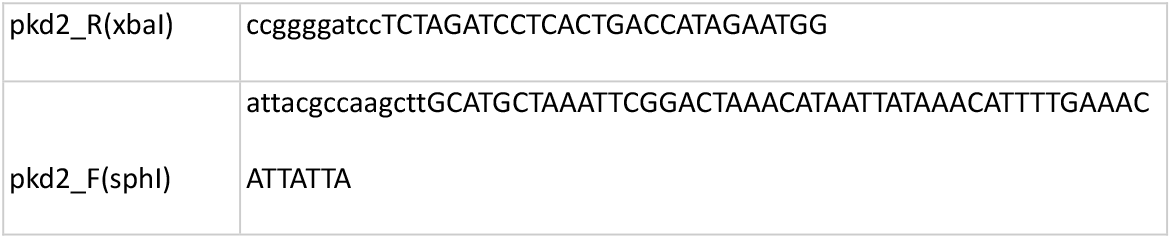

Germ-line transformation was performed using standard microinjection techniques (Mello et al. 1991). Pkd-2prom::3xnovoGFP-CLA-1, pkd-2prom::BFP, pCFJ90, and PPD49.26 were co-injected into the germline.

### Microscopy and image analysis

Animals were anesthetized with 20 mM levamisole and mounted on their back on 5% agar pads on glass slides. We used 1-day-old males to observe RnB axon morphology. To observe anterior-posterior and circumferential axons, animals were observed with fluorescence microscopy (Zeiss Axio Imager A1) with 100x magnification. To observe diagonal commissures, we utilized Nikon CSU-W1spinning disk confocal microscopy with 100x magnification.

### RnB synapse imaging

To visualize the presynaptic pattern of RnB neurons, we utilized EM1554 bxIs30[pkd-2^prom^:: gfp; pkd-2^prom^::RAB-3-Mcherry; ttx-3p:: GFP] II; him-5(e1490)V and EM1890 bxIs33[pkd-2^prom^::3xnovoGFP-CLA1, pkd-2p::RAB-3-Mcherry, pkd-2p::BFP] I ; him-5(e1490)V. Animals were anesthetized with 20 mM levamisole and mounted on their back on 5% agar pads on glass slides. Images were obtained with Nikon CSU-W1spinning disk confocal microscopy with 100x magnification with 5 nm thickness. All images were obtained with same exposure time for the same wave length. (For EM1554, 488 nm laser power 15%, exposure time 50 ms, 561 nm laser power 25%, exposure time 500 ms. For EM1855, 405 nm laser power 30%, exposure time 600 ms, 488 nm laser power 20%, exposure time 200 ms.)

To quantify the protein levels of pre-synaptic markers, all planes for each animal were z-stacked. The circle ROI, or region of interest, contained 205 pixels and was set on the synaptic ring, followed by measuring mean signal intensity. We also quantified the signal intensity of the cytoplasmic fluorescent markers in the same ROI and calculated pre-synapse/cytoplasmic signal intensity ratio.

## Results

### Reporter genes used in this study

Two integrated transgenes were used to visualize the trajectories and synapses of ray B-type sensory neurons (Fig 2*A*). Both utilized the promoter of the polycystin-homolog gene *pkd-2* (Barr and Sternberg, 1999) to drive expression of fluorescent markers in the RnB neurons. In addition to the RnB’s, this promoter drives expression in tail sensory neuron HOB and four CEM sensory neurons in the male head (Barr and Sternberg, 1999).

**Fig. 2.**
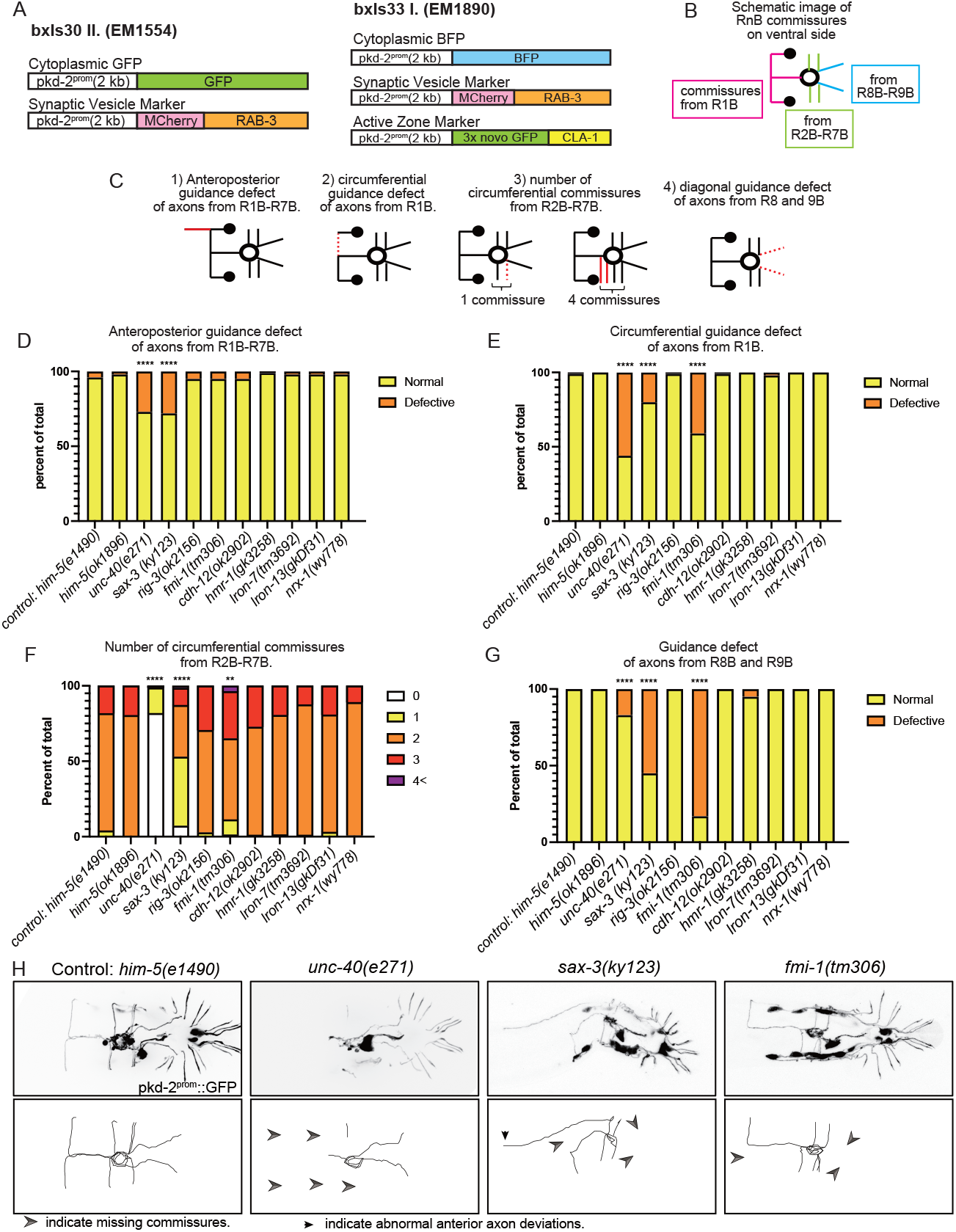
Trajectories of RnB neurons in CAM mutants. (A) Transgenes labeling axon trajectories and pre-synapses in EM1554 *bxIs30*[*pkd-2*^*prom*^::GFP, *pkd-2*^*prom*^::MCherry-RAB3, *ttx-3*^*prom*^::GFP] II ; *him-5(e1490)*V and in EM1890 *bxIs33*[*pkd-2*^*prom*^::BFP, *pkd-2*^*prom*^::MCherry-RAB3, *pkd-2*^*prom*^::3xnovoGFP-CLA-1, *ttx-3*^*prom*^::MCherry] I ; *him-5(e1490)*V. (B) A diagram showing typical RnB commissures in wild type. (C) Diagrams showing a typical example of each anatomical defect. Red lines indicate the commissure with defect. (D-G) Quantified data for defect of anterior-posterior axons (D), of circumferential axon guidance (E), in number of circumferential commissures from R2-7B (F), and of posterior commissures (G). ****p<0.0001 vs him-5, **p<0.01 vs him-5, fisher’s exact test. (H) Fluorescent images of RnB neurons and HOB neurons (top panels) and traces of commissures from Ray neurons (bottom panels) of *him-5(e1490)* control animals and mutant animals. RnB neurons as well as HOB neuron were marked with *pkd-2*^*prom*^::GFP. Arrow heads indicate the missing commissures and arrows indicate the abnormal anterior axon deviation.

For visualization of neuron trajectories, both transgenes included Ppkd-2 driving a cytoplasmic marker. One transgene, *bxIs30II* (strain EM1554), expressed cytoplasmic GFP along with mCherry-labeled Rab-3, the *C. elegans* homolog of mammalian Rab3, a component of synaptic vesicles (Nonet et al., 1997; Lázaro-Peña et al., 2018). The second, *bxIs33I* (strain EM1890), included cytoplasmic BFP along with RAB-3-mCherry and 3xnovoGFP-CLA1. CLA-1 is a piccolo/bassoon-like scaffolding protein active zone component (Xuan et al., 2017). Both strains carried a mutation in the *him-5* gene, resulting in the presence of males in selfing hermaphrodite populations.

### Axon trajectory phenotypes of strains carrying single mutations in CAM genes

The trajectories of the RnB neuron processes in wild type, visualized by the cytoplasmic marker, are shown in Fig 1*A* and schematically in Fig 2*B*. Based on the characteristics of RnB axon morphology, we classified the trajectory defects in mutants into one of four categories (Fig. 2*C*): 1) anteroposterior guidance defect of axons from R1B-R7B. 2) circumferential guidance defect of axons from R1B. 3) number of circumferential commissures from R2B-R7B. 4) posterior commissures of axons from R8 and 9B. We quantified the percentage of defective worms in each of those four categories (Fig. 2*D-G*).

The CAM genes and mutations studied in this work are shown in Table 1. We determined the effect of mutations in all of the B-neuron-expressed genes except *pan-1*, which is embryonic lethal. We observed trajectory defects in three of the genes tested, *unc-40, sax-3*, and *fmi-1. unc-40* mutants had significant abnormalities in the dorsoventral guidance system for formation of all the ray commissures (Fig. 2*E, F, H*), as reported previously (Hedgecock et al., 1990; Jia and Emmons, 2006). Significant defects were also observed in anteroposterior guidance and posterior commissures (Fig, 2*D, G*). Although posterior commissures were reported to be unaffected in a previous paper (Jia and Emmons, 2006), some commissures were missing when we observed with a confocal microscope. *sax-3* mutants had abnormal anteroposterior and circumferential axonal projections (Fig. 2*D-F*). The abnormal anterior axonal projection was often seen in combination with a deletion of the circumferential axon or R1B (Fig. 2*H*). The lack of circumferential R1B axon seems to be due to a failure of turning of the anterior-posterior axon from the cell body. In *sax-3* mutants, the posterior commissures from rays 8 and 9 were also frequently missing (Fig 2*G, H*). Mutation in *fmi-1*/flamingo resulted in a deficit of the circumferential commissures (Fig.2*E, F, H*), and most notably, an absence of the posterior commissures in almost all animals (Fig.2*G, H*). For all of the remaining genes, *rig-3, cdh-12, hmr-1, lron-7, lron-13* and *nrx-1*, animals containing single mutations had no significant defects in axon trajectories when compared to wild type (Fig.2*D-G*).

### Axon trajectory phenotypes of strains carrying multiple mutations in CAM genes

In view of the lack of effects of single mutations in most of the genes, we asked whether this was due to redundancy of their activities by examining double and multiple mutants. We found that this does not appear to be the case. Since *rig-3* and *lron-7* loci are close and we could not create this double, we created two quintuple mutants (*hmr-1*; *cdh-12 lron-13*; *nrx-1*; *lron-7* and *hmr-1*; *cdh-12 lron-13*; *nrx-1*; *rig-3*). In all of the strains carrying multiple mutations in genes with no single effect, there were no significant abnormalities (Fig. 3*A-D*).

**Fig. 3.**
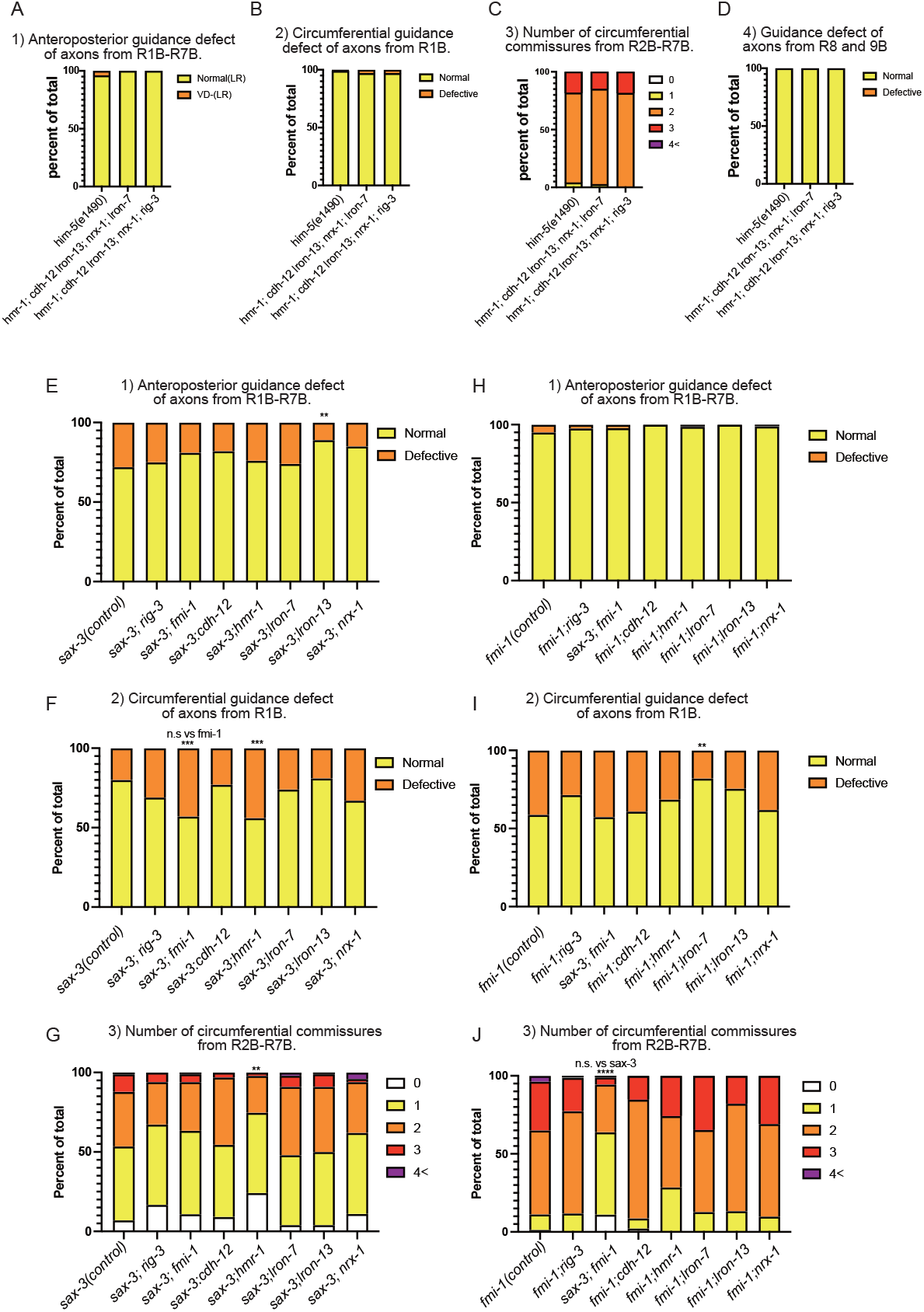
Genetic interactions between CAMs expressed in RnB neurons. (A-D) Percentage of defect of anterior-posterior axon guidance (A), percentage of defect of circumferential R1B axon guidance (B), number of circumferential commissures from R2-7B (C), and percentage of defect of posterior commissures (D) for quintuple mutants of CAMs expressed in RnB neurons. (E-F) Percentage of defects of anterior-posterior axon guidance (E), percentage of defects of circumferential R1B axon guidance (F), and number of circumferential commissures from R2-7B (G) for double mutants of *sax-3* and CAMs expressed in RnB neurons. (H-J) Percentage of defective anterior-posterior axon guidance (H), percentage of defective circumferential R1B axon guidance (I), and number of circumferential commissures from R2-7B (J) for double mutants of *fmi-1* and CAMs expressed in RnB neurons. ****p<0.0001 vs control, *** p<0.001 vs control, **p<0.01 vs control, *p<0.05 vs control, n.s p>0.05, fisher’s exact test.

Next, we crossed the strains with observable phenotypes, *sax-3* and *fmi-1*, into the strains with no obvious phenotype to look for genetic interactions (enhancement or suppression) (Fig. *E-J*). We found that the *hmr-1* mutant significantly enhanced the axon guidance defects in the circumferential commissures in *sax-3* mutant (Fig. 3*F, G*), indicating that *hmr-1* and *sax-3* have redundant functions. In addition, *lron-13* and *lron-7* suppressed axon guidance defects in *sax-3* and *fmi-1*, respectively (Fig. 3*E, I*). These mutants have not been reported to have any significant phenotypes, but they may be involved in the regulation of axon guidance in cooperation with other CAMs.

### Effects of mutations in genes expressed in RnA neurons on axon trajectories of RnB neurons

RnA and RnB neuron processes run side by side from the dendrites in the rays through circumferential commissure fascicles until they enter the pre-anal ganglion (Fig 1*B*, Fig 2*B*). We therefore examined whether CAMs expressed in RnA neurons affect axon guidance in RnB neurons. Although many cell adhesion molecules are expressed in the RnA neurons (Fig. 1*C*), we focused on the effects of four genes (*casy-1, bam-2, sax-7*, and *tol-1*) that are widely expressed in most of the RnA neurons. Single mutations in *casy-1, bam-2*, and *sax-7* showed no abnormalities in the morphology of the RnB neurons (Fig.4*A-D*). In the *tol-1* mutant, there was a significant increase in the number of commissures from Ray2-7 (Fig. 4*C*). This result suggests that *tol-1* may be involved in the fasciculation of the circumferential commissures.

**Fig. 4.**
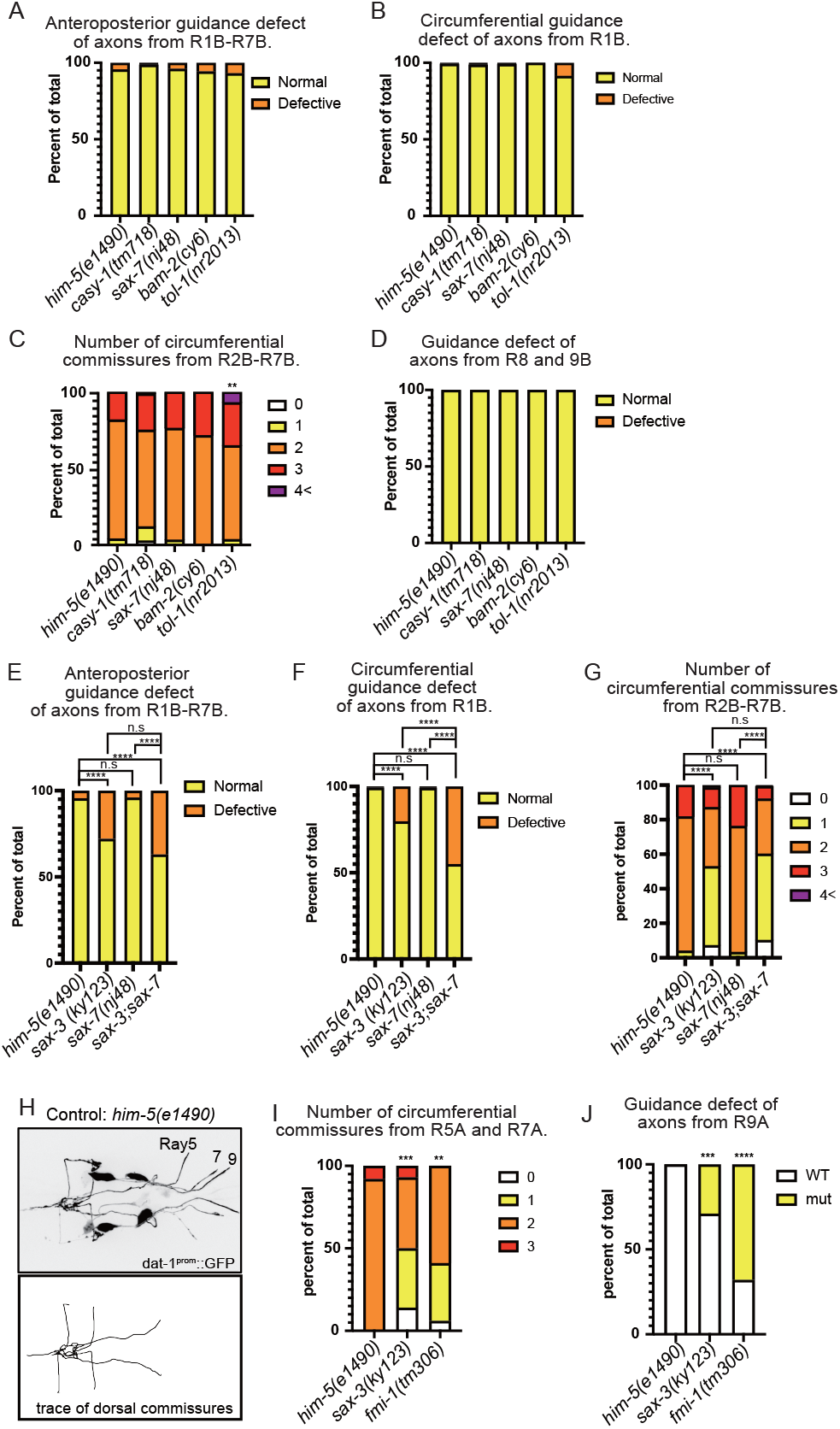
*sax-3* and *fmi-1* mutants showed abnormal RnA neuron morphology. (A-D) Percentage of defective anterior-posterior axon guidance (A), percentage of defective circumferential R1B axon guidance (B), number of circumferential commissures from R2-7B (C), and percentage of defective diagonal commissures for mutants of CAMs expressed in RnA neurons (D). (E-G)Percentage of defective anterior-posterior axon guidance (E), percentage of defective circumferential R1B axon guidance (F), and number of circumferential commissures from R2-7B (G) for *sax-3, sax-7*, and *sax-3; sax-7* mutants are shown. (H) Fluorescent images of R5A, R7A, R9A neurons (top panels) and traces of commissures from Ray neurons (bottom panels) of *him-5(e1490)* control animal. RnA neurons were marked with *dat-1*^*prom*^::GFP. (I-J) Percentage of number of commissures from R5, 7A(I) and percentage of defective diagonal commissures (J) for sax-3 and fmi-1 mutants are shown. ****p<0.0001 vs him-5, *** p<0.001 vs him-5, **p<0.01 vs him-5, Fisher’s exact test.

SAX-7 has been reported to interact with SAX-3 in axonal fasciculation (Chen et al., 2019). We therefore asked whether these proteins acted together in a single pathway for ray fasciculation or separately by determining whether mutation in *sax-7*, expressed in A, would enhance the defect of RnB axon trajectory in a double mutant with *sax-3*, expressed in B (Fig.4*E-G*). We found the *sax-7* mutation significantly enhanced the B-type abnormalities in the circumferential commissure of the *sax-3* mutant (Fig. *4F*). This suggests that these two proteins have independent and apparently somewhat redundant functions in promoting fasciculation of their respective neuron processes in the commissures.

In view of this possibility, we tested for a possible reciprocal effect on RnA trajectories of mutations in those RnB-expressed genes that affected RnB trajectories. We visualized a subset of RnA neurons using DAT-1p::GFP expressed in R5A, R7A, and R9A (Fig. 4*H*). We observed abnormalities in RnA neurite trajectories in both *sax-3* and *fmi-1* mutants similar to those observed in the RnB neurons (Fig. 4*I, J*). Thus, each of the two neuron types appears to promote the trajectories of the other type, consistent with their co-fasciculation in the outgrowing commissures.

### Effects of mutations in CAM genes on synapse formation

CAMs are involved not only in axon guidance but also in synapse formation (Shen and Bargmann, 2003; Dalva et al., 2007; Kim and Emmons, 2017; Kurshan et al., 2018; Lázaro-Peña et al., 2018; Philbrook et al., 2018). We visualized RnB synapses with fluorescent markers localized respectively to synaptic vesicles (RAB-3) and pre-synaptic densities (CLA-1). Both reporter genes showed a pattern of fluorescence consistent with RnB pre-synapses as described by electron microscopy (Jarrell et al., 2012) (Fig 5*A, B*). Most RnB synapses are in the pre-anal ganglion, where the densest fluorescence appears as a ring, dubbed here the “synaptic ring.” In addition, consistent with the mixed axonal/dendritic character of most *C. elegans* neurons, they receive a similar amount of input from other neurons. These post-synapses are not visualized here. Multiple pre-synapses were also observed within the pre-anal ganglion along the midline axon and in the lumbar ganglia, but few in circumferential commissures and, in the case of the RAB-3 reporter, almost none in the rays themselves. By contrast, the GFP::CLA-1 reporter indicated the presence of presynaptic densities in the rays (Fig. 5*B*). We examined EM images of the rays and found synapse-like structures with high electron density within the ray dendrites in the rays (Fig. 5*C*). Altogether, in the EM reconstruction of a single adult male, the RnB neurons form 1143 chemical synapses onto 123 other sensory neurons, including other ray neurons, interneurons, and motor neurons, most of which are in the pre-anal ganglion (Jarrell et al., 2012) (Fig. 1B).

**Fig. 5.**
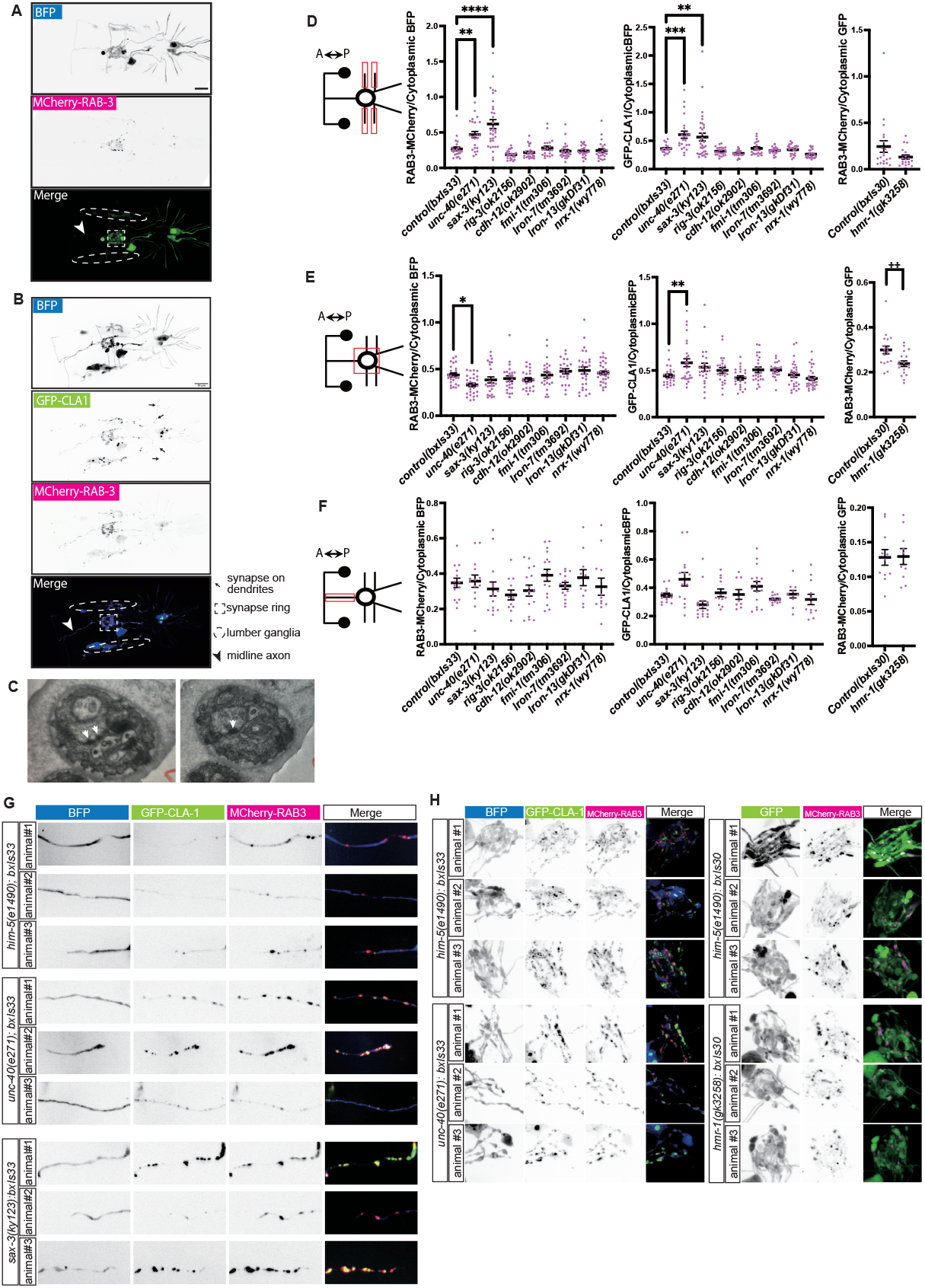
Expression of synaptic markers in CAM gene mutants. (A) Fluorescent images of EM1554. Broken-line square, broken-line oval, and arrowhead represent synapse ring, lumbar ganglia, and midline axon, respectively. (B) Fluorescent images of EM1890. Black arrow, broken-line square, broken-line oval, and arrow represent synapses on ray dandrites, synaptic ring, lumber ganglia, and midline axon, respectively. (C) Electron microscopy images of two consecutive sections. The white arrows indicate synapse-like structures with high electron density. (D-F)Schematic images and quantified data of pre-synaptic markers in (D)Circumferential commissures, (E) Synaptic ring, and (F) midline axon. Data of left panels show signal intensity ratio of 3xnovoGFP-CLA1/cytoplasmic BFP for each mutant with *bxIs33*. Center panels show signal intensity ratio of RAB3-MCherry/cytoplasmic BFP for each mutant with *bxIs33*. Right panels show signal intensity ratio of RAB3-MCherry /cytoplasmic GFP for each mutant with *bxIs30*. Red boxes in the schematic images indicate the region of interest. ****p<0.0001 vs control, *** p<0.001 vs control, **p<0.01 vs control, multiple comparison test. ++ p<0.01 vs control, t-test. (G) Fluorescent images of circumferential commissures in mutants with *bxIs33* or *bxIs30*. (H) Fluorescent images of synaptic ring in mutants with *bxIs33*.

To assay the amount of synapse formation, in order to control for different levels of gene expression we determined the ratio of signal intensity between the synaptic signals (RAB-3::mCherry for *bxIs30II*, RAB-3::mCherry and GFP::CLA-1 for *bxIs33I*) and the cytoplasmic signals (GFP for *bxIs30II*., BFP for *bxIs33I*)(Fig. 5D-F). In *unc-40* and *sax-3* mutants, both pre-synaptic marker ratios were increased in circumferential commissures (Fig. 5D and G). This suggests accumulation of synaptic proteins in processes that have failed to reach their targets. In the *unc-40* mutant, the RAB-3-MCherry ratio decreased in the synaptic ring region but the GFP-CLA1 ratio increased (Fig. 5E, H). Thus CLA-1 protein may accumulate within those few processes that reach the target area. In the *hmr-1(gk3258)* mutant, the RAB-3::mCherry signal ratio was decreased in the synaptic ring, whereas there was no significant increase or decrease within circumferential commissures (Fig. 5D-F). (The CLA-1 reporter could not be tested because of proximity of the mutation to the transgene insertion site.) Thus *hmr-1* may be involved in accumulation of synaptic vesicles. Besides these, there were no other deviations from wild type ratios in the single and multiple mutants. In none of the mutants was there any significant pre-synapse signal ratio change in midline axons (Fig. 5F).

These results indicate a limited effect of mutations in RnB-expressed CAM genes on accumulation of pre-synaptic components within the neuronal processes. Whether this indicates these CAM genes play no role in recognition of synaptic targets and formation of synapses or that synaptic components accumulate in the absence of target recognition and synapse formation cannot be determined.

### SAX-3 regulates circumferential axon guidance in slt-1 independent manner

For two of the genes we studied, *sax-3* and *fmi-1*, interacting partners or pathway components are known in other systems. For SAX-3/ROBO, it has been demonstrated in many species that Slit, a secreted LRR protein, functions as a ligand (Brose et al., 1999; Zallen et al., 1999; Hao et al., 2001; Dickinson and Duncan, 2010). In our expression dataset, *slt-1* is mainly expressed in ALN neurons and dorsal body wall muscles. We tested whether *slt-1* is involved in RnB axon guidance and found that *slt-1* null mutants showed abnormal anterior axon projections and loss of circumferential axons, which were also observed in *sax-3* mutants (Fig. 6*A, B*). The absence of *slt-1* did not enhance the *sax-3* mutant phenotype in a double mutant, consistent with SLT-1 functioning as a ligand for SAX-3. By contrast, *slt-1* single mutation did not show any significant abnormality in the number of commissures from R2-7B (Fig. 6*C*). This result and the lesser overall severity of the *slt-1* null mutant phenotype compared to the *sax-3* null mutant phenotype indicate that molecules other than SLT-1 may function as ligands for SAX-3 in RnB axon guidance.

**Fig. 6.**
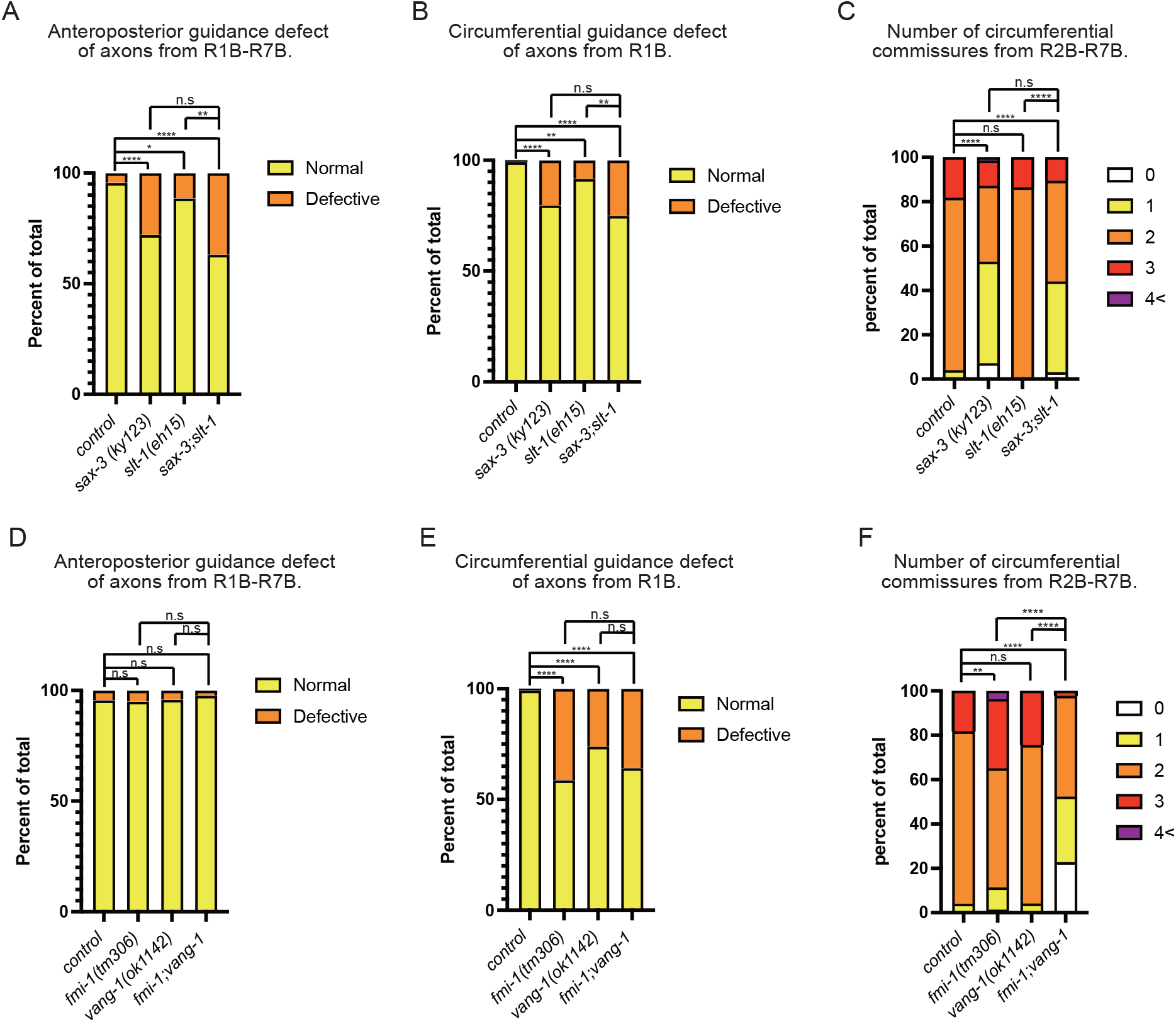
Genetic interactions between *sax-3, fmi-1*, and their known interacting partners. (A-C) Percentage of defective anterior-posterior axon guidance (A), percentage of defective circumferential R1B axon guidance (B), and number of circumferential commissures from R2-7B (C) for *sax-3, slt-1*, and *sax-3; slt-1* mutants. (D-F) Percentage of defective anterior-posterior axon guidance (D), percentage of defective circumferential R1B axon guidance (E), and number of circumferential commissures from R2-7B(F) for *fmi-1, vang-1, and fmi-1; vang-1* mutants. ****p<0.0001, **p<0.01, *p<0.05, n.s p>0.05, Fisher’s exact test.

### FMI-1 regulates RnB axon guidance both in PCP pathway and PCP independent pathway

In Drosophila, Flamingo, a homologue of *fmi-1*, was previously reported to work together with Van Goh and Frizzled to regulate the PCP (planar cell polarity) pathway (Usui et al. 1999). We tested whether the PCP pathway genes are involved in RnB axon guidance (Fig. 6*D-F*). The lack of the circumferential axon from R1B was occasionally seen in a *vang-1* mutant (homolog of Van Goh) (Fig. 6*G*). This phenotype was not enhanced by *fmi-1* null mutant (Fig. 6*D*), indicating *fmi-1* and *vang-1* work in the same pathway. On the other hand, the *fmi-1; vang-1* double mutant showed a significantly more severe phenotype than the *fmi-1* mutant alone in circumferential commissures from R2-7 (Fig. 6*F*). These results suggest that *fmi-1* and *vang-1* regulate the morphology of RnB neurons within in the same pathway as well as in parallel pathways.

## Discussion

*C. elegans* ray sensory neurons provide an excellent opportunity to identify the factors that determine their wiring selectivity. Genes involved in formation of the two neuron types and that specify ray neuron identities have been described, including transcription factors and signaling pathways (Baird et al., 1991; Chow and Emmons, 1994). Regulatory genes must act by controlling the expression of effector genes that function during morphogenesis. Here we examined the class of genes thought to specify the cell surface labels that govern cell-cell recognition for synapse formation. Like other neurons in the *C. elegans* nervous system, ray sensory neurons express multiple cell adhesion molecules (CAMs), including many conserved across species (Hahn and Emmons, 2003; Taylor et al., 2021; this laboratory unpublished). Reflecting an aspect of ray identity, the combination expressed by the neurons in each ray pair differs (Fig 1*C*). The same genes are expressed in other combinations across the entire set of neurons and muscles in the posterior nervous system, indeed, across the entire animal in all tissues, reflecting their presumptive function as neutral cell labels (Taylor et al., 2021; this laboratory unpublished).

Our primary result reported here is that mutations in the CAM genes expressed by the ray neurons have surprisingly limited effects on two aspects of ray neuron phenotype: axon trajectories and accumulation of pre-synaptic proteins in axonal processes. General absence of effects on axon trajectories reflects the robustness of the mechanisms that guide formation of the structure. For example, we found that SAX-3/ROBO and SAX-7/L1CAM each appear to independently contribute to fasciculation of the RnB and RnA circumferential processes.

Lack of effects on accumulation of synaptic proteins probably reflects the fact that assembly and transport of such components to sites of expected synapse formation may be to some degree independent of target recognition and synapse formation (Kurshan and Shen, 2019). Methods such as iBlinc or GRASP that more directly assay the presence of a connection between pre- and post-synaptic cells can be used to seek further evidence of synapse formation (Feinberg et al., 2008; Desbois et al., 2015).

Still, we observed increased synaptic proteins in circumferential commissures in some mutants. The accumulation of synapse proteins was examined in three different compartments. the kinetics of synaptic proteins at the three locations did not coincide. For example, in the sax-3 mutant, synapse protein accumulation was increased in the circumferential commissure, but not in the synapse ring and longitudinal commissure (Fig. 6D-F). This may suggest that the degree of contribution of CAMs to synaptogenesis varies from compartment to compartment within the same nerve.

We observed CLA-1-GFP puncta in dendritic region in RnBs. We also observed synapse-like structures in electron microscopic images. Functional dendritic spines have been reported in the GABAergic motor neurons of *C. elegans* (Cuentas-Condori et al., 2019). The accumulation of CLA-1-GFP in male *C. elegans* ray nerves can be other examples of functional dendritic synapses.

## Data availability

Strains and plasmids are available upon request. The datasets and imaging data generated and/or analysed during the current study are available from the corresponding author on reasonable request.

## Acknowledgement

We thank H Bülow and D Hall for technical advice. The plasmid containing CLA-1-GFP was kindly gifted by P Kurshan. *C. elegans* strains were provided by the CGC, which is funded by NIH Office of Research Infrastructure Programs (P40 0D010440), and NBRP (Japan).

This work was supported by NIH grants from NIHD (P30HD071593 to S.W.E.), NIMH (R01MH112689 to S.W.E.), NIGMS (R01GM066897 to S.W.E.), JSPS Overseas Research Fellowships to N.S., and the Analytical Imaging Facility of the Albert Einstein Cancer Center (P30 CA013330).

---

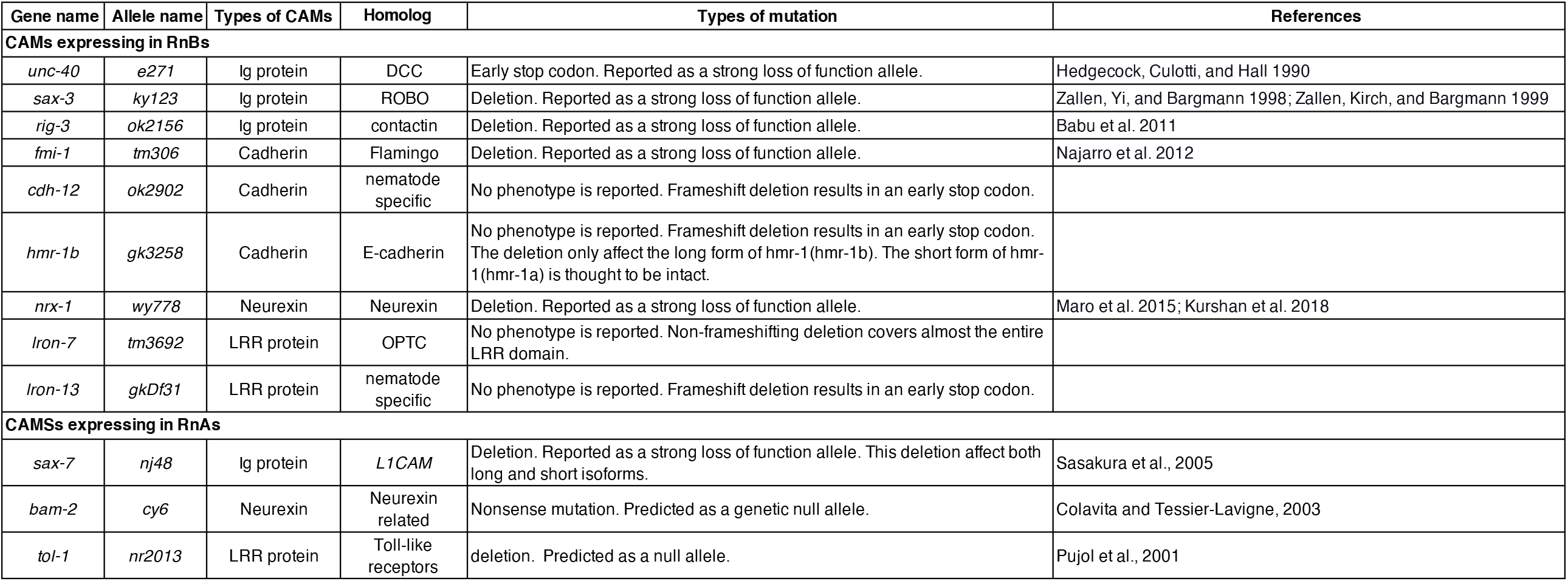

